# Democratizing DNA Fingerprinting

**DOI:** 10.1101/061556

**Authors:** Sophie Zaaijer, Assaf Gordon, Robert Piccone, Daniel Speyer, Yaniv Erlich

## Abstract

We report a rapid, inexpensive, and portable strategy to re-identify human DNA using the MinION, a miniature sequencing sensor by Oxford Nanopore Technologies. Our strategy requires only 10-30 minutes of MinION sequencing, works with low input DNA, and enables familial searches. We also show that it can re-identify individuals from Direct-to-Consumer genomic datasets that are publicly available. We discuss potential forensic applications as well as the legal and ethical implications of a democratized DNA fingerprinting strategy available to the public.

---

DNA is a powerful biometric identifier. With the exception of monozygotic twins, DNA profiles are unique for each individual on earth, leave traces that can stay for an extended period of time after the person of interest is gone, and enable the identification of individuals that are not part of the reference database *via* familial searches (*1–3*). Current DNA forensic techniques generate profiles mainly by genotyping a small set of autosomal polymorphic short tandem repeats (STRs) (*4*). This method usually requires transporting samples to labs equipped with bulky equipment such as capillary electrophoresis machines and thermocyclers. As of today, the state of the art forensic genotyping platforms (e.g. DNAscan or RapidHIT 200) take about 90 minutes to process a DNA sample, weigh over 50 kilograms, have a capital cost of more than a quarter of a million dollars, and require about $300 to process a sample (*5*). These technological barriers limit DNA fingerprinting to high-latency applications and narrow its accessibility to specific societal entities such as law enforcement or security forces.

Here, we demonstrate a portable and inexpensive strategy for re-identification of anonymous human DNA using a MinION sequencer (Oxford Nanopore Technologies). MinION is a radically different DNA sequencer compared to other contemporary platforms (*6, 7*): it weighs less than 100 grams, costs $1000, and only requires a standard laptop to operate. The data is available in near real-time, with the first sequenced DNA fragments ready for analysis only a few seconds after the start of a sequencing run. Due to its small footprint and high robustness, it is possible to employ MinION sequencers outside of laboratories. Early access users have generated MinION sequencing data in various places, including rural Africa (*8, 9*), hotel rooms, and classrooms (*10*). The company recently announced efforts to further miniaturize the sequencer as a cellphone adaptor and revealed plans to use the device as an autonomous sensor that enables Internet of Things (IoT) applications (*11*).

MinION technology poses several challenges for human re-identification. The primary challenge is the high sequencing error rate, as about 10-25% of the reported bases are erroneous compared to <0.5% by a mainstream Illumina platform (*12, 13*). The error profile is dominated by insertions and deletions (indels), which complicate precise measurements of STR length polymorphisms. The MinION sequencing throughput is also many orders of magnitudes smaller than other massively parallel sequencing platforms, yielding only 3-50 million bases per run (equal to ~0.01 × coverage of a human genome).

We developed a robust strategy for human DNA re-identification, termed the ‘MinION sketch’ (*14*). The sketching strategy rapidly generates extremely low coverage shotgun sequencing data using the MinION sequencer and then compares this data to a reference database. Our method has no PCR step to eliminate the latency introduced by DNA amplification and to increase portability. However, the extremely low coverage of MinION data limits the ability to interrogate genomic sites multiple times, resulting in low variant calling confidence and the inability to distinguish homozygous from heterozygous variants. To address these challenges, we developed a Bayesian algorithm that computes a posterior probability that the sketch matches an entry in the reference database (**Fig. S1**). The algorithm takes into account the error rate in MinION data, the inability to observe both haplotypes, the allele frequency, and the prior probability of a match. It works in a sequential manner and updates the posterior probability with every new read that is available until a match is found.

MinION sketching provides sufficient data for human DNA identification even after a few minutes of operation. As an initial experiment, we generated a sketch from Craig Venter’s genomic DNA (Coriell’s GM25430) and applied the Bayesian algorithm to match the sketch to his genome-wide genotyping array (*14*), which is publicly available (*15*). After analyzing 13 minutes of MinION sketching, the *posterior* probability of a positive match surpassed 99.9% (**Fig. S2A**). Despite the high error rate and sparsity of the genotyping array, the algorithm needed only 155 bi-allelic variants to reach this conclusion, only ~2 times above the theoretical expectation to re-identify a person by fingerprinting random bi-allelic markers (*16*). The initial selection of the prior probability had no effect on the matching ability of the method and only slightly increased the time required to achieve a high confidence match. With a prior probability that considers a database ten times bigger than the world’s population (10^−11^), the posterior probability reached 99.9% after 18 minutes of sketching (**Fig. S2B**). As a negative control for our algorithm, we compared Craig Venter’s sketch to a genome-wide array of an unrelated random individual. The matching algorithm reported a probability of less than 10^−5^ that Craig Venter’s sketch matches this person.

Next, we tested MinION sketching to specifically find a person in a much larger collection of real genomic data (**Fig. 1A**). To this end, we compiled a collection of 19,000 genome-wide genotyping arrays from individuals that were tested with Direct to Consumer (DTC) genetic testing companies such as 23andMe, AncestryDNA, and Family Tree DNA (*14*). Then, we generated a MinION sketch from the saliva of a male individual (YE001) whose 23andMe data are publicly available and were part of the collection of 19,000 individuals. The Bayesian algorithm blindly searched the 19,000 collection and identified YE001 with a posterior probability greater than 99.9% with only 14 minutes of sketching. None of the other 19,000 samples reached this level of probability of identification, indicating that specificity is achieved even for larger databases.

We explored DNA re-identification with the MinION using low input DNA and portable equipment. For DNA collection, we again obtained a saliva sample from YE001 but this time we used an FTA card that yielded 50ng of DNA. Next, we prepared the MinION library with BentoLab (*17*), a consumer product recently funded by a Kickstarter campaign to provide a simple portable laboratory (**Fig. S3A**). The pre-release version of this device weighs ~3kg and includes a heating plate and a small centrifuge with the footprint of a small laptop. We processed the sample with BentoLab and minimal wet-lab equipment and prepared a 1D MinION library that relies on single adapter ligation (*14*). The 1D protocol requires a lower amount of input DNA but results in much noisier data since each DNA molecule is read once as opposed to twice as in the previous experiments. Next, we ran this sketch against the collection of 19,000 genomes. Despite all of these limitations, thirty minutes of sketching was sufficient to reach over a 99% matching probability to the YE001 sample and to differentiate it from the other 19,000 genomes (**Fig. S3B,C**).

Finally, we found that MinION sketches also enable familial searches to identity first-degree relatives (**Fig. S4A**). We modified the Bayesian algorithm to consider first-degree matches in addition to the exact matches (*14*). We then generated a MinION sketch for a 1000 Genomes sample (NA12890) and compared the sketch to her son’s whole genome (NA12977). After 25min of sketching, the Bayesian framework reported her son as a first-degree match with a probability greater than 99.9% (**Fig. 1B**). As expected, the exact match posterior was well below 10^−5^ almost the entire time (**Fig. S4B**). We also compared the NA12890 sketch to her own whole genome sequencing data. The algorithm correctly reported her genome as an exact match and not as a first-degree match (**Fig. S4B**). To further challenge our method, we also matched the sketch to her granddaughter’s genome (NA12879). Again, the posterior for a direct match was nearly zero for all tested conditions (**Fig. S4B**); the posterior for a first-degree match had a brief 1 minute spike towards 100% due to one long read but quickly decayed and rejected the hypothesis that the sample is a first degree relative (**Fig. S4C**). Finally, we also calculated the probability of familial matching between the NA12890 sketch and the 19,000 DTC genomes. As expected, no match was found, highlighting the specificity of the method in identifying first-degree relatives (**Fig. S4D**).

To summarize, our experiments show that a few minutes of MinION sequencing provides sufficient information for DNA re-identification. Our method works with either whole genome reference data or common DTC genotyping datasets whose generation is highly cost-effective (low hundreds of dollars per sample) and within the same price range as the generation of forensic profiles such as the CODIS or ENFSI sets. Even with this proof of principle, the sample weight for MinION sequencing can go as low as 50ng without challenging the protocol to a great extent and the method is capable of first-degree familial searches. ONT recently launched a new chemistry design (R9) for the MinION sequencer that ratchets bases through the nanopore five times faster than the version used for these experiments. If successful, this chemistry will provide sufficient data for identification in less than five minutes of sequencing, getting closer to real time person identification.

Our strategy highlights the possibility of using the MinION in combination with other off-the-shelf equipment to build an inexpensive system for low latency DNA fingerprinting. This may open new possibilities for security and law enforcement applications. Near-real-time DNA surveillance can serve as a tool for identification of victims after a mass disaster or for border control to fight human trafficking. These two applications can benefit from the familial search capability of the sketch that enables families to donate their own DNA to identify their loved ones. Portable fingerprinting by MinION sketching can also offer great advantages for regular forensic casework, but integration might be more challenging. Existing forensic databases only hold STR profiles of individuals that are not compatible with our MinION sketches. However, with the continuous drop in costs for sequencing and genotyping arrays, developing combatable databases might be more economically feasible in the future. From a legal perspective, forensic use of MinION sketching seems to comply with US federal and state statutes. Previous scholarly work has postulated that genotyping DNA of abandoned material by law enforcement is lawful in all US states (*18*), permitting the generation of MinION sketches from such material by forensic teams. A more complicated aspect is the legality of constructing genome-wide genotyping databases by compulsory DNA collection from arrestees, similar to current forensic databases. Indeed, genome-wide genotyping would collect far more information than the small number of STR markers that are currently used. However, the US legal framework mostly places restrictions on the purpose of compulsory DNA collection rather than the extent of genotyping (*19*). In a major decision, the Supreme Court of the United States recently stated that DNA collection from arrestees indeed constitutes a search under the 4th Amendment; but if DNA is solely used for identification, this procedure is reasonable and viewed as a sophisticated version of standard arrestee booking procedures such as fingerprint collection (*20*). The Court further stated that genotyping markers that are associated with disease predisposition is not a significant invasion of privacy as long as they are not tested for that end. Therefore, it seems that a genome-wide forensic database would not violate the 4^th^ Amendment.

In addition to forensics applications, MinION sketches could be used in the future by citizen scientists (or biohackers) to fingerprint anonymous samples for self-curiosity, *sous*veillance (*21*), or activism. The equipment used for our experiments is relatively inexpensive, portable, mostly accessible for a lay-audience and not substantially complicated to operate. Indeed, most of the MinION sequencing data presented here was generated by inexperienced students as part of a computer science class (*10*). In addition, our results show that MinION sketches are fully compatible with data from DTC genomic companies. Several websites, such as OpenSNP.org (*22*) and the Personal Genome Project (*23, 24*), publicly share large amounts of identifiable genomic data from DTC participants. The current information in these websites theoretically enable MinION re-identification of tens of thousands individuals using direct and familial searches.

While our results show that re-identification is technically feasible, MinION sketching might be illegal when used outside of law enforcement and security applications. For instance, France and New York State prohibit DNA genotyping without the consent of the sample provider (*25*). In Israel, the Genetic Information Law, it is illegal to conduct DNA tests without accreditation to operate a genetic lab (*26*). In the US, a company recently had to pay over two million dollars to two employees after conducting a simple DNA fingerprinting test that violated the Genetic Information Non-discrimination Act (GINA) (*27*). With the increased interest in this sequencer as a consumer device, educational tool, or autonomous IoT sensor, the MinION sequencer will reach new users from a vast range of fields that are not fully aware for these nuances. Therefore, it might also be advisable to consider the development of mitigation steps to avoid misusage by users and regulation complications. Mitigation could in part rely on technical solutions imposed by the manufacturer; these can range from real-time notification for users when human DNA is registered by the cloud-based base-caller, to the real-time rejection of DNA molecules mapped to the human genome (e.g. using the method described by (*28*)) unless the user is authorized for analyzing human DNA. Beyond that, the community could start considering “sequencing etiquette” and expected norms for mobile sequencing. All of these would be highly important to facilitate the transition of this exciting technology from the research community to the general public.

**Figure 1.**
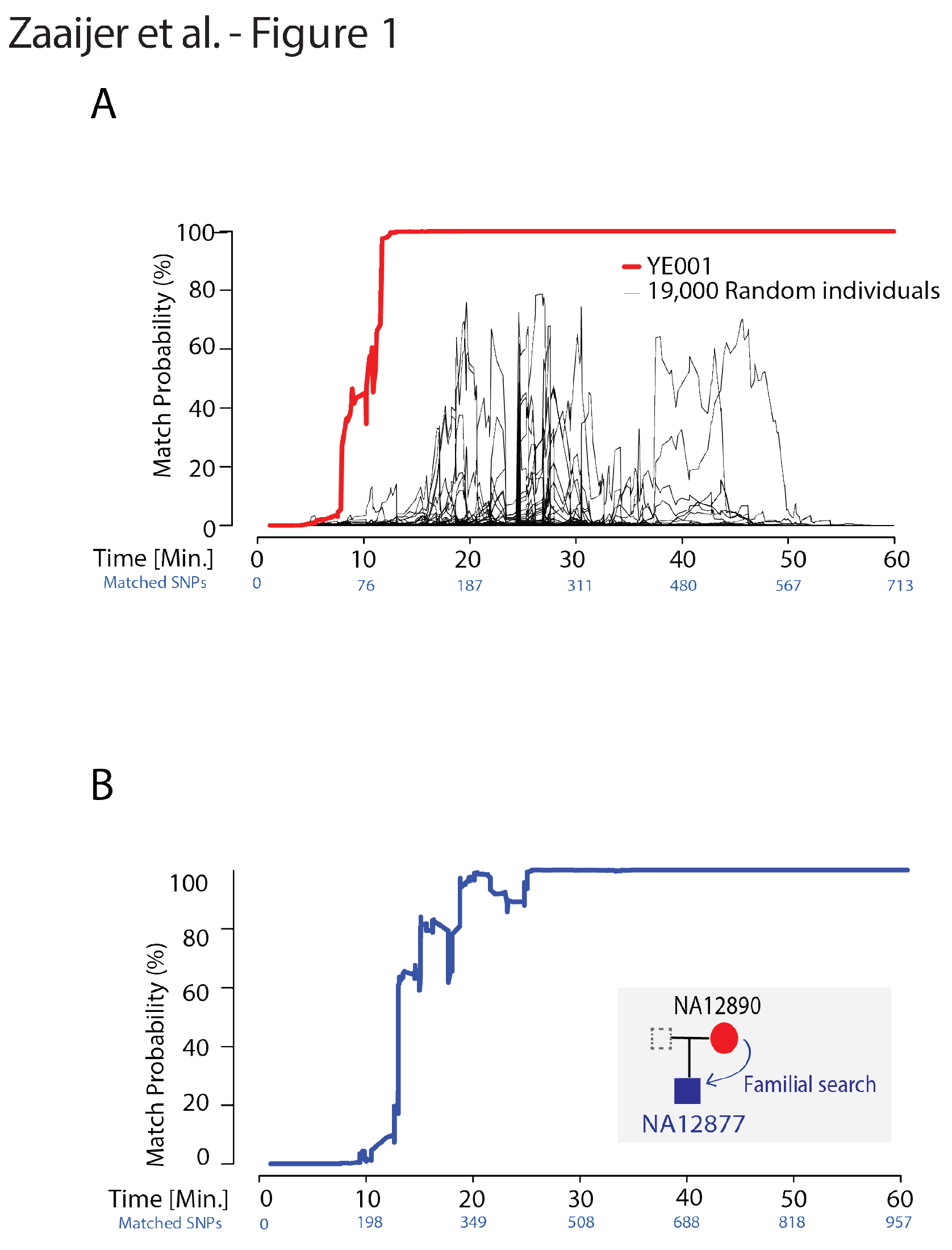
Exploiting MinION sequencing for human re-identification. The figure shows the posterior probability of a match as a function of MinION sketching time. Matched SNPs denote the number of SNPs used for the posterior computation by the algorithm in each time point (A) Searching a collection of 19,000 DTC genome-wide arrays with YE001’s MinION sketch. Red: posterior probability for an exact match to the correct genome-wide array; black: posterior probability for random individuals in the collection (B) An example of a familial search. Blue: the posterior probability for a familial match between the sketch of NA12890 and her son’s reference genome.

## Acknowledgments

Y.E. holds a Career Award at the Scientific Interface from the Burroughs Wellcome Fund. This study was supported by a generous gift from Andria and Paul Heafy (Y.E.) and National Institute of Justice (NIJ) grant 2014-DN-BX-K089. We thank D. Zielinski and T. Willems for useful comments, W. Stephenson and K. Pandit from New York Genome Center’s Innovation lab for technical assistance, B. Wolfenden, P. Boeing, E. Jorgensen for early access to the Bento Lab, M. Micorescu from Oxford Nanopore Technologies for useful discussions, O. Tene and O. Yakobovich that familiarized us with the concepts of sousveillance, and the Columbia Ubiquitous Genomics class 2015 for data generation.

## References and Notes

1. M. Gymrek, A. L. McGuire, D. Golan, E. Halperin, Y. Erlich, Identifying personal genomes by surname inference. Science 339, 321 (2013).

2. M. Kayser, P. de Knijff, Improving human forensics through advances in genetics, genomics and molecular biology. Nature Reviews Genetics 12, 179 (2011).

3. F. R. Bieber, C. H. Brenner, D. Lazer, Finding criminals through DNA of their relatives. Science, 5778, 1315 (2006).

4. C. Smith, S. Strauss, L. DeFrancesco, DNA goes to court. Nat Biotechnol 30, 1047 (2012).

5. L. K. Hennessy et al., Developmental validation studies on the RapidHIT™ human DNA identification system. Forensic Science International: Genetics Supplement Series 4, e7 (2013).

6. N. J. Loman, M. Watson, Successful test launch for nanopore sequencing. Nature methods 12, 303 (2015).

7. Y. Erlich, A vision for ubiquitous sequencing. Genome research 25, 1411 (2015).

8. J. Quick et al., Real-time, portable genome sequencing for Ebola surveillance. Nature 530, 228 (2016).

9. T. Hoenen et al., Nanopore sequencing as a rapidly deployable Ebola outbreak tool. Emerging infectious diseases 22, 331 (2016).

10. S. Zaaijer, Y. Erlich, Using mobile sequencers in an academic classroom. Elife 5, e14258 (2016).

11. E. Young, in The Atlantic. (2016).

12. C. L. Ip et al., MinION Analysis and Reference Consortium: Phase 1 data release and analysis. F1000Research 4, (2015).

13. M. Jain et al., Improved data analysis for the MinION nanopore sequencer. Nature methods 12, 351 (2015).

14. See supplementary materials.

15. S. Levy et al., The diploid genome sequence of an individual human. PLoSBiol 5, e254 (2007).

16. Z. Lin, A. B. Owen, R. B. Altman, Genomic research and human subject privacy. Science 305, 183 (2004).

17. Bento Lab. Nat Biotech 34, 455 (05//print, 2016).

18. E. E. Joh, Reclaiming’Abandoned’DNA: The Fourth Amendment and Genetic Privacy. Northwestern University Law Review 100, 857 (2006).

19. B.-J. Koops, M. Schellekens, Forensic DNA phenotyping: regulatory issues. Colum. Sci. & Tech. L. Rev. 9, 158 (2008).

20. Maryland v. King, US Supreme Court, 2013, vol. 1958.

21. S. Mann, J. Nolan, B. Wellman, Sousveillance: Inventing and Using Wearable Computing Devices for Data Collection in Surveillance Environments. Surveillance & Society 1, 331 (2002).

22. B. Greshake, P. E. Bayer, H. Rausch, J. Reda, OpenSNP-a crowdsourced web resource for personal genomics. PLoS One 9, e89204 (2014).

23. M. P. Ball et al., A public resource facilitating clinical use of genomes. Proceedings of the National Academy of Sciences 109, 11920 (2012).

24. L. Sweeney, A. Abu, J. Winn, Identifying participants in the personal genome project by name. Available at SSRN 2257732, (2013).

25. S. Soini, Genetic testing legislation in Western Europe—a fluctuating regulatory target. Journal of community genetics 3, 143 (2012).

26. J. Zlotogora, Genetics and genomic medicine in Israel. Molecular genetics & genomic medicine 2, 85 (2014).

27. S. Miller, United States District Court for the Northern District of Georgia Finds Employer Liable for Violation of Genetic Information Nondiscrimination Act (” GINA“) in the Case of the” Devious Defecator“-Lowe v. Atlas Logistics Group Retail Services, LLC 1. American Journal of Law and Medicine 41, 684 (2015).

28. M. Loose, S. Malla, M. Stout, Real time selective sequencing using nanopore technology. bioRxiv, 038760 (2016).

